# Gut microbiome transition across a lifestyle gradient in Himalaya

**DOI:** 10.1101/253450

**Authors:** Aashish R. Jha, Emily R. Davenport, Yoshina Gautam, Dinesh Bhandari, Sarmila Tandukar, Katharine Ng, Susan Holmes, Guru Prasad Gautam, Jeevan Bahadur Sherchand, Carlos D. Bustamante, Justin L. Sonnenburg

## Abstract

The composition of the gut microbiome in industrialized populations differs from those living traditional lifestyles. However, it has been difficult to separate the contributions of human genetic and geographic factors from lifestyle/modernization. Here, we characterize the stool bacterial composition of four Himalayan populations to investigate how the gut community changes in response to shifts in human lifestyles. These groups led seminomadic hunting-gathering lifestyles until transitioning to varying dependence upon farming. The Tharu began farming 250-300 years ago, the Raute and Raji transitioned 30-40 years ago, and the Chepang retain many aspects of a foraging lifestyle. We assess the contributions of dietary and environmental factors on their gut microbiota and find that the gut microbiome composition is significantly associated with lifestyle. The Chepang foragers harbor elevated abundance of taxa associated with foragers around the world. Conversely, the gut microbiomes of populations that have transitioned to farming are more similar to those of Americans, with agricultural dependence and several associated lifestyle and environmental factors correlating with the extent of microbiome divergence from the foraging population. For example, our results show that drinking water source and solid cooking fuel are significantly associated with the gut microbiome. Despite the pronounced differences in gut bacterial composition across populations, we found little differences in alpha diversity across populations. These findings in genetically similar populations living in the same geographical region establish the key role of lifestyle in determining human gut microbiome composition and point to the next challenging steps of isolating dietary effects from other factors that change during modernization.

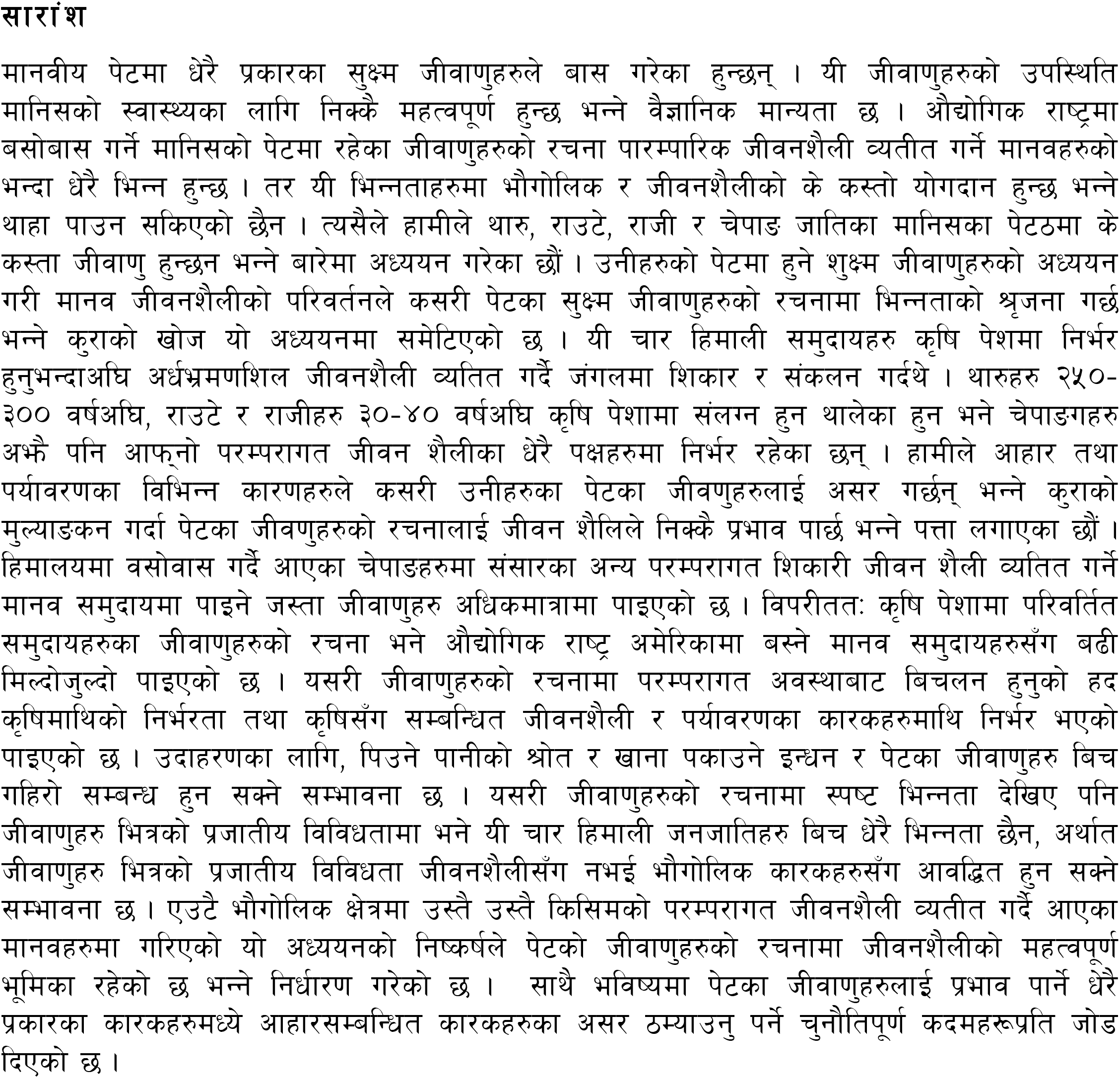

## INTRODUCTION

The human gut is comprised of a diverse community of bacteria, the microbiome or microbiota, that influences several aspects of human physiology including nutrient metabolism, immune responses, and resistance to infectious pathogens [1–3]. This highly malleable microbial component of human biology exhibits rapid, and in some cases, irreversible changes in response to dietary and environmental factors [4–11]. Modern humans experienced diverse environments since expanding out of Africa ~100,000 years ago, and over the past ~10,000 years hunting and gathering has largely yielded to different forms of agriculturally supported lifestyles. Dietary changes combined with a variety of other factors associated with the industrial revolution have been credited as contributing to the alterations in the gut microbiome in industrialized populations [12]. However interpretation of the current data is clouded by potential contributions of human genetic variation, environment, and geographical factors [5,7,13]. The potential connection between the gut ecosystem and several chronic diseases necessitates a better understanding of the extent to which modernization has contributed to population-wide community changes during industrialization [14,15].

Comparisons of the gut microbiomes of traditional human populations in Africa and South America with those of the industrialized Western populations from Europe and USA reveal that the human gut microbiome varies across geography and lifestyles [16–30]. One universal trend from these studies is the higher diversity of gut bacteria in unindustrialized traditional populations. However, most of the traditional societies investigated thus far live within tropical latitudes [31]. Hence, whether difference in alpha diversity is due to contrasting lifestyles, residence in the tropics, or other factors remains unclear. Moreover, most studies to date compare the gut microbiomes between populations that reside in geographically distinct regions, represent extreme modes of human subsistence, and are genetically and culturally distinct [16,17,19–21,25,27]. Although some studies have attempted to mitigate these differences by comparing human populations that reside in close geographical proximities [23,24,28], these populations have been separated for tens of thousands of years, a period of time sufficient for genetic and cultural differences to arise [32]. Since gut microbiome can be influenced by genetic, environmental, and cultural factors [23,28,33], these variables make it difficult to determine the impact of lifestyle changes in the gut microbiome in such distinct populations. Hence, understanding how transitions in human lifestyles lead to changes in the gut microbiomes would be greatly aided by studying populations that have undergone recent changes in culture, lifestyle, and diet.

In order to explore how the gut microbiota changes as human populations transition from traditional to more urban lifestyles, we have analyzed the gut microbiomes from four rural Himalayan populations and compared them to those of Americans with European ancestry. The Himalayan populations include the Chepang – a foraging population, the Raute and Raji – two foraging communities that are currently transitioning to subsistence farming, and the Tharu – former jungle dwellers that have completely transitioned to farming within the last two centuries. We assessed contributions of lifestyle, diet, and environment on the gut microbial variation in the rural Himalayan populations. Our results show that gut microbiome composition mirrors the transitions from traditional to westernized lifestyle in Himalaya. In addition to the dietary gradient across these populations, intra- and inter-population variability in lifestyle elucidated additional environmental and lifestyle associations that may contribute to microbiota change.

## RESULTS

### Description of populations

Our participants included 54 individuals from four Himalayan groups, including Chepang (N=14), Raji (N=9), Raute (N=11), and Tharu (N=20) with median age of 40 years (SD ± 14 years) from rural villages in Nepal (Figure 1, Supplementary Table 1). These four populations are long-term residents of the Himalayan foothills (altitude less than 1000 m) and contain various degrees of East Asian ancestries [34–36]. Although all four of the Himalayan populations in this study were forest dwellers until recently [37–40], habitat loss due to rapid deforestation, population expansions of non-native groups, establishment of new settlements, and construction of modern highways led to settlement of these groups at various time points in the last 300 years.

**Figure 1:**
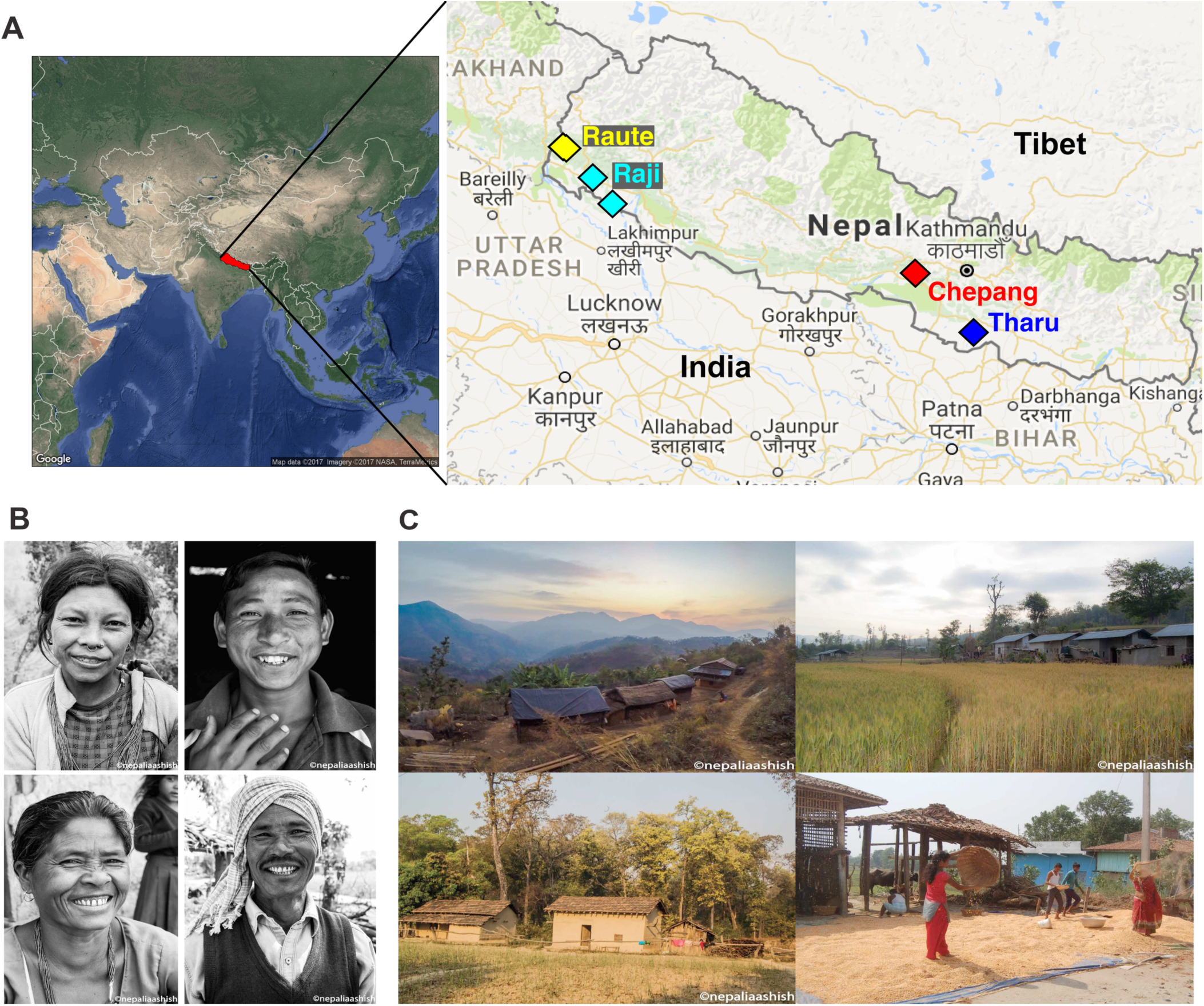
Sampling locations and habitats of the Himalayan populations in Nepal. **(A)** Map displaying the geographical locations of sampled villages in southern Nepal (altitudes<1000 meters above the sea level; latitude 26.97-29.15). The Tharu are geographically most distant from the Raute and Raji and reside closer to the Chepang. **(B)** Individuals representing each population. From top-left in clockwise direction: a Chepang female, a Raute male, a Tharu male, and a Raji female. **(C)** Habitats of each population. From top-left in clockwise direction: the remote Chepang village, Raute village, Tharu harvesting rice, and Raji village.

Historical records indicate that the Tharu gradually transitioned into agrarian lifestyles beginning in the late eighteenth century (250-300 years ago) [40]. They have fully transitioned into farming and are virtually completely disengaged from foraging practices. Historically, the Raute, Raji, and Chepang were semi-nomadic foragers and their diets included native tubers, greens and fruits from the jungle, wild honey, fish, and occasional game [38,39,41]. The Raute and Raji abandoned their foraging lifestyles in the 1980s [37,38]. While the Raute have settled in the remote hills in Far-Western Nepal, the Raji have settled in the Terai plains, which is relatively more urbanized. The Chepang were fully nomadic at least until 1848 [42] and began supplementing their foraging practices with subsistence agriculture less than a century ago [39]. The Chepang in this study currently inhabit a remote village that is devoid of modernity, including electricity, running water, irrigation, fertilizers, modern machines, and marketplaces. They still practice slash and burn agriculture and are completely dependent on rainwater for farming. Because yields from such traditional farming are low, their daily diet consists of wild plants such as *sisnu* (nettles) that are foraged from the forests.

### Lifestyle gradients in the Himalayan populations

We conducted surveys to assess how lifestyle changed as these seminomadic populations transitioned to farming in the last few hundred years. The survey questionnaire included questions pertaining to current dietary practices, traditional and modern medicines, and several environmental factors, including sources of drinking water, alcohol use, and tobacco consumption (N=53, Supplementary Table 2). Combustion of solid biomass fuel such as firewood or animal dungs produces environmental particulate matters increasing indoor air pollution [43]. Prolonged exposure to environmental pollutants has the potential to alter gut microbiome [44]. Hence, we assessed the fuel types used for cooking and location of kitchen in our Himalayan participants. We also surveyed presence of parasites in our participants microscopically.

Supervised learning using a Random Forest classifier (RFC) model on the survey data (including intestinal parasite load) assigned the individuals to their respective populations with high accuracies (100% accuracy for the Chepang and Tharu, 90% for the Raute and Raji, OOB error = 3.5%, Supplementary Table 3A), indicating these populations have distinct lifestyles. A correspondence analysis (CA) of the survey data (including intestinal parasite load) also revealed lifestyle differences between these populations (Figure 2A). The first CA dimension (CA1) explained 15.8% variation in the data and was strongly correlated with lifestyle gradients. Along CA1, samples progressed from the Chepang foragers at one extreme, to the Raute and Raji transitioning populations, and then to the Tharu farmers at the opposite extreme (Figure 2B). Despite the geographical distance between them, the Raji lifestyle appears to be more similar to that of the Tharu farmers, consistent with the Raji settlement occurring in a more urbanized setting compared to the Raute. Similarly, the Raute reside in geographical proximity to the Raji, although their lifestyle partitions between the Raji and the Chepang, indicating geographical proximity is not driving the lifestyle differences.

**Figure 2:**
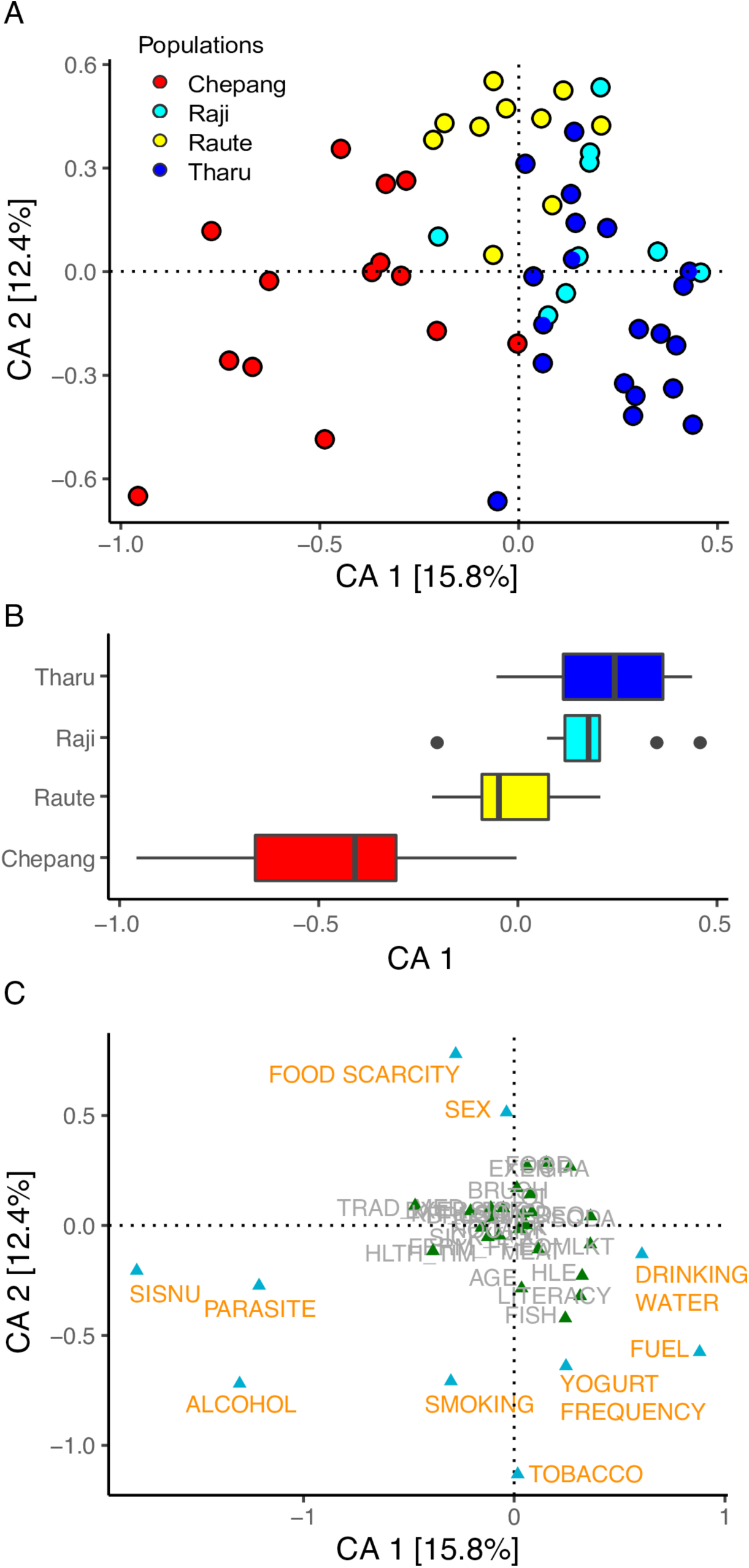
Correspondence Analysis based on survey questionnaires and parasite assessment in the Himalayan populations. First two dimensions of the correspondence analysis and the amount of variation explained are shown. **(A)** Each circle represents an individual and colors represent the populations. **(B)** Distribution of populations along the primary CA1 axis shows patterns of separation by lifestyles. Chepang foragers (red) and Tharu farmers (blue) are on two extreme ends of CA1. In between the two are the Raute (yellow) and Raji (cyan), the two communities that are transitioning from foraging to farming. **(C)** Factors in gold are those that have more than expected eigenvalues and thus contribute most to the top two dimensions in the Correspondence Analysis.

A total of 10 variables contributed highly to the first two CA dimensions and most of them are strongly associated with dietary differences and modernity (Figure 2C). These differences are described in details in Supplementary Figure 1. Briefly, foraged plants such as *sisnu* (nettles) and *jaand*, a slushy alcoholic beverage made from fermenting millet or corn, are staples of the Chepang diet. In contrast, *sisnu* and *jaand* consumption was minimal among the Raute, Raji, and Tharu. Also, perceived food scarcity was higher in the Chepang and Raute relative to Raji and Tharu. Although meat consumption was low across all four populations, the Tharu consumed animal products such as yogurt more frequently than the other three populations. Furthermore, the Tharu and Raji also showed increased signs of modernity. For example, they have installed tube wells at their homes, enabling access to underground water for drinking. In contrast, the Chepang and Raute still fetch drinking waters from rivers and streams. Also, use of solid biomass fuel was lower in Tharu and Raji while Chepang and Raute are still completely dependent on burning firewood for cooking. Although we detected low overall levels of intestinal parasites across the participants, Ascaris, Entamoeba, Trichuris, Hymenolepis, and Coccidia were detected in some, and most of the infected were the Chepang. Together, the diet and lifestyle assessments provide unbiased support that the four populations represent a gradient from traditional to increasingly agrarian and urban lifestyles.

### Gut microbiome composition varies by lifestyles

In order to assess whether the gut microbiome varies across lifestyles, we characterized the gut bacterial composition of these populations using the Illumina MiSeq to sequence the V4 region of 16S ribosomal RNA (rRNA) gene obtained from a total of 79 stool samples (including technical replicates) with an average of 11,570 (±4653) high quality reads/sample (Supplementary Figure 2, Supplementary Table 4). Since flash freezing of the samples was not possible in the remote sampling areas in the Himalaya, we used commercially available DNAgenotek OMNIgene kits to collect stool samples from the four populations (N=54). We also collected stool samples from 10 Americans of European descent using OMNIgene kits and compared them with freshly frozen samples to evaluate whether preservation method affected microbiome profile. The 16S rRNA profiles of the same samples stored by flash freezing or by OMNIgene were remarkably similar, with reproducible differences in minor taxa (Euryarcheota and Cyanobacteria), demonstrating the reliable preservation of microbiome composition with the OMNIgene kits (Supplementary Figure 3). Due to the reproducible, albeit minor, differences between the two collection methods, we used the OMNIgene data from the Americans for consistency in subsequent comparative analyses.

Comparison of the community structure in the five study populations using unweighted UniFrac distances, a measure of compositional similarity that includes the phylogenetic relatedness between microbiomes, showed that the gut microbial composition varied across populations (P< 2.2 X 10^-16^, *Kruskal-Wallis test*). The four Himalayan populations exhibited much larger distances when compared to the Americans than when compared to one another (Supplementary Table 5). The Chepang were the most distant from the Americans followed by the Raute, while the Raji and Tharu were equally close to the Americans. Within Himalaya, the Chepang were more distant from the Tharu and Raji relative to the Raute while the Raute, Raji and Tharu were equally distant from one another. Similar results were also observed with weighted UniFrac and Bray-Curtis distances, both of which take the taxa abundance into account (Supplementary Table 5).

Visualization of these distances using a Principal Coordinates Analysis (PCoA) revealed separation of populations along the top two dimensions (p=1 × 10^-5^, *PERMANOVA*, Figure 3A). Furthermore, gradients in lifestyles were reflected by the distribution of populations along the primary axis (PCoA1, Figure 3B). These distributions remained consistent when using Bray-Curtis and weighted UniFrac distances as well (p=1 × 10^-5^ for both, *PERMANOVA*, Supplementary Figure 4 and 5). When American microbiomes were eliminated from the principal coordinate analyses, the gradient between the Himalayan populations remained pronounced (P=1 × 10^-5^, *PERMANOVA*, Supplementary Figure 6). Among the four Himalayan populations, the strongest separation was observed between the Chepang foragers and the Tharu farmers.

**Figure 3:**
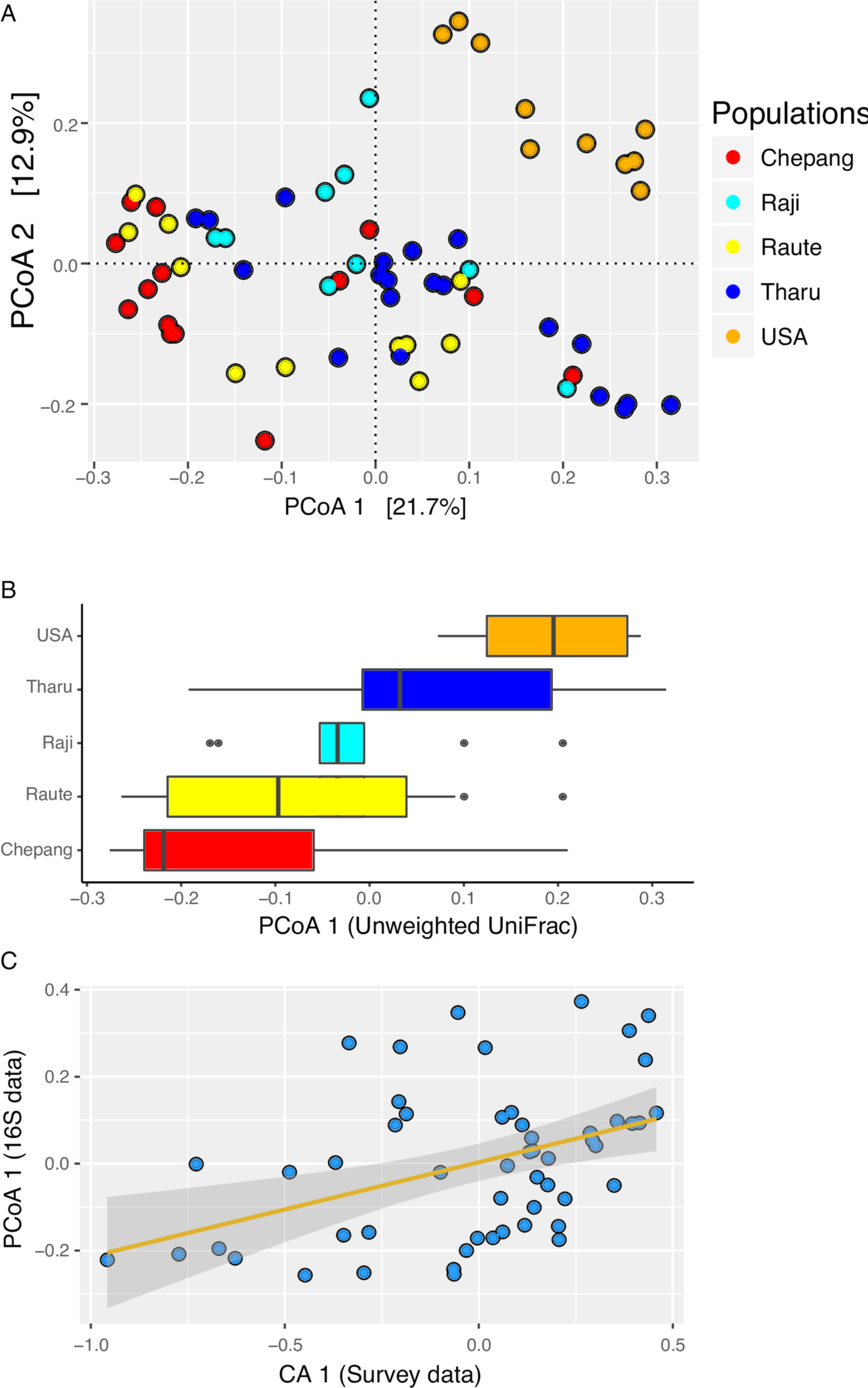
Gut microbiome compositions show gradients in lifestyles. **(A)** PCoA of the unweighted UniFrac distances colored by populations. Each dot represents an individual and colors indicate the populations. Chepang foragers (red), Raute (yellow) and Raji (cyan) communities that are transitioning from foraging to farming, Tharu farmers (blue), and Americans (orange). **(B)** Distributions of populations along the PCoA1 axis show patterns of separation by lifestyles. **(C)** Gut microbial composition of the Himalayan populations represented by the primary dimension of the unweighted UniFrac distance (PCoA1) strongly correlates with lifestyle differences represented by the top dimension of the corresponding analysis performed on the survey data (CA1, Spearman’s rho = 0.44 and P-value = 0.001). Correlation between CA2 and PCoA1 was not statistically significant.

A random forest classifier based on the 16S rRNA-defined read sequence variant (16S RSV) data assigned the Chepang, Tharu, and American individuals to their respective source populations with 86%, 100%, 100% accuracies (OOB error=32%, Supplementary Table 3B). The classification accuracy for the Raute and Raji, the two populations that recently transitioned from foraging to farming, were relatively poor (<10%). While some of the individuals from these groups were classified as the Chepang, others were classified as the Tharu. However, none of the Himalayan individuals were classified as American. These results collectively show that the gut microbiome compositions of the Himalayan populations are distinct from those of the Americans. They also indicate that within Himalaya, the gut microbiome of the Chepang foragers differs from that of the Tharu farmers while that of the Raute and Raji reflect their transitional state in their lifestyles.

To formally evaluate whether variation in gut microbiota reflects lifestyle differences within Himalaya, we assessed associations between the respective primary dimensions from the lifestyle questionnaire and parasite analysis (CA1) and gut microbial composition analysis (PCoA1) (Figure 3C and Supplementary Figure 4). We found that the CA1 was strongly correlated with the PCoA1 obtained from all of the three distance matrices (Spearman’s rho = 0.47, 0.44, and 0.28 for Bray-Curtis, unweighted UniFrac, and weighted UniFrac distances, respectively, P-value < 0.05 for all three, *correlation test*). The CA1 was also correlated with PCoA2 of all three distance matrices (Spearman’s rho = 0.26, 0.44, and 0.39; P-value = 0.06, 0.001, and 0.004 for Bray-Curtis, unweighted UniFrac, and weighted UniFrac distances, *correlation test*). Conversely, no significant correlations were detected between CA2 and either of the PCoA axes from all three distances (P-value < 0.05, *correlation test*). Notably, CA1 but not CA2 is associated with lifestyle gradient (Figure 2). Strong and consistent correlations between CA1 and PCoA axes indicate that gut microbiome compositions of the Himalayan populations mirror their lifestyles.

### Gut bacterial diversity (alpha diversity) does not vary across lifestyles

Previous studies have suggested that elevated species diversity in gut microbiome is a hallmark of traditional populations [19,28]. We assessed the alpha diversity in the five study populations using four measures, namely species richness, Fisher’s alpha, Shannon’s H, and Simpson’s D at various rarefaction depths ranging from 500-3000 reads (Figure 4). Species richness and Fisher’s alpha were not significantly different between any of the five populations (Bonferroni adjusted P>0.05, *Kruskal-Wallis test*). We did find marginally significant differences in Shannon and Simpson indices between these populations (Bonferroni adjusted P<0.05, *Kruskal-Wallis test*). A post-hoc pairwise comparison of all five populations showed that only the alpha diversity in the Tharu was slightly lower than that in the Americans (Bonferroni adjusted P = 0.02 and 0.03 respectively, *Dunn’s test*) and none of the of the four diversity measures showed significant differences in alpha diversity between the Chepang, Raute, Raji, and the Americans. Moreover, correlations between each of the four alpha diversity measures and lifestyle differences within Himalaya measured using the CA1 were not statistically significant (P>0.05, *correlation test*). These results indicate that lifestyle differences among the Himalayan populations or between these populations and Americans have little effect on the alpha diversity of the gut microbiome.

**Figure 4:**
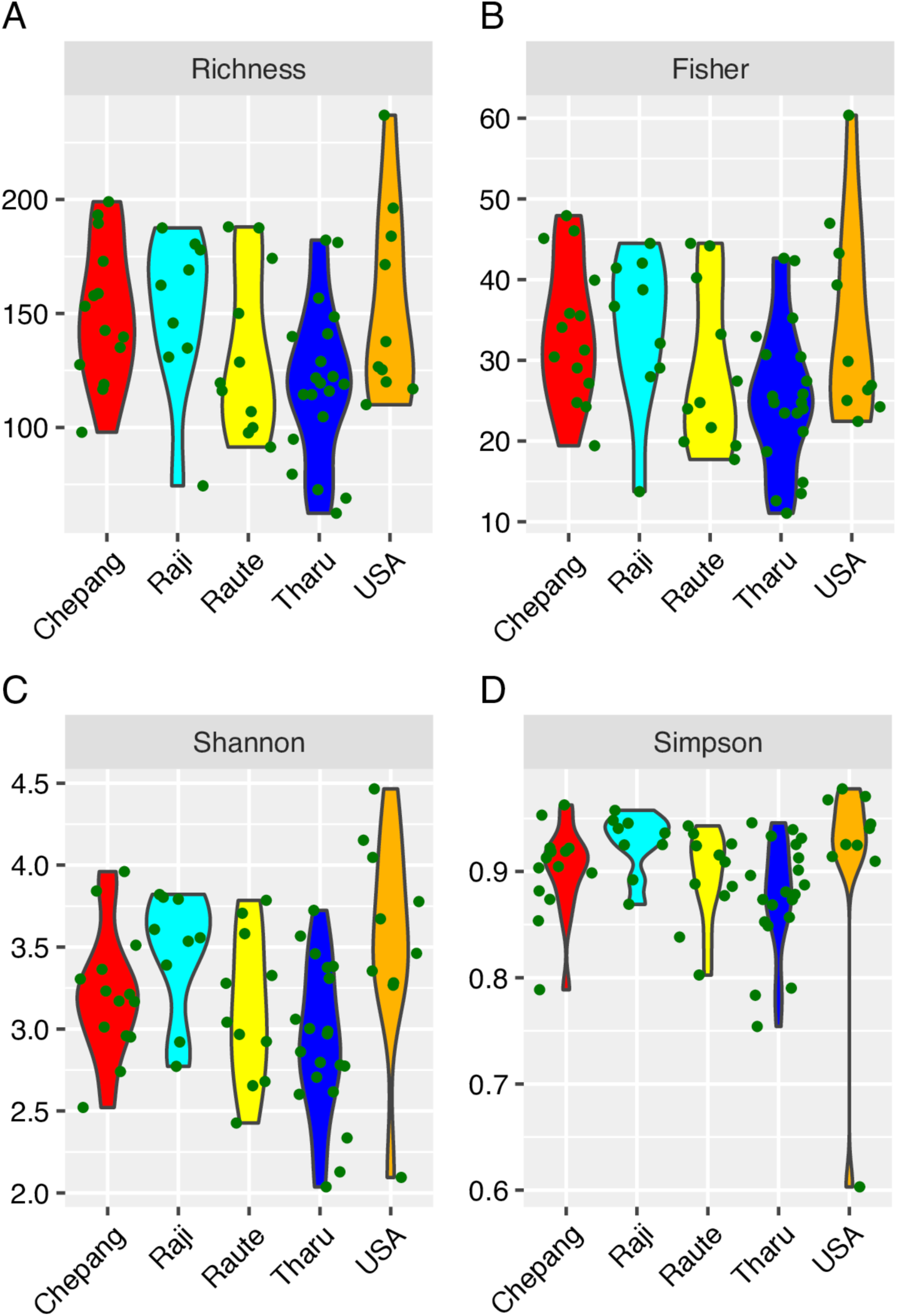
Alpha diversity in the study populations. Alpha Diversity calculated at a rarefaction depth of 3000 reads per sample. No significant differences in species richness (A) and Fisher’s alpha (B) were detected between the five study populations. Shannon’s H (C) and Simpson’s D (D) were significantly lower in the Tharu relative to the Americans. No differences in any of these four

### Bacterial taxa are associated with lifestyle transitions

Although lifestyle differences have little effect on the alpha diversity, gut microbiome compositions of the Himalayan populations reflected the gradients in their lifestyles. To identify taxa driving the differentiation of the gut microbiomes across lifestyles we compared the differences in abundance of individual phylum across the five populations using a negative binomial generalized linear model (GLM) as implemented in DESeq2 [45]. Differential abundances were detected for 6 out of 10 phyla (FDR adjusted P-value <0.05, *GLM,* Supplementary Table 6) and four of the six phyla reflect a traditional-western lifestyle gradient. The Himalayan populations were characterized by higher abundance of Proteobacteria, while abundances of Actinobacteria, Firmicutes, and Verrucomicrobia were highest in the Americans, intermediate in the farmers (Tharu, Raji, and Raute), and lowest in the Chepang foragers (Figure 5A). Higher levels of Proteobacteria and lower levels of Actinobacteria and Verrucomicrobia are common features of many traditional human gut microbiomes across the world [19,24,28,30].

**Figure 5.**
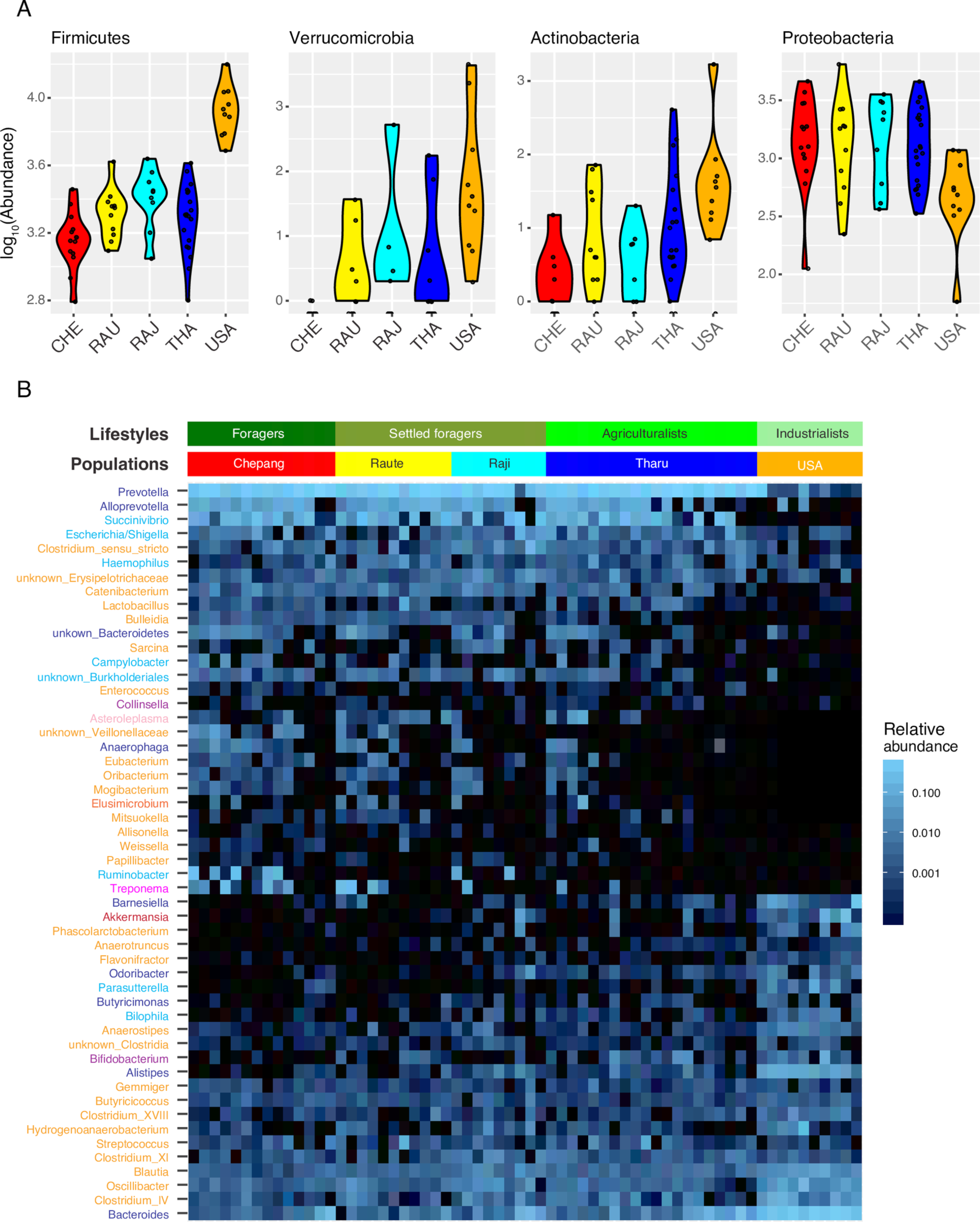
Distinctions in the gut microbiome across lifestyles. **(A)** Phyla with most significant differences in abundances between the five populations. Abundances of Firmicutes, Verrucomicrobia, and Actinobacteria reflect gradients of traditional-western lifestyles. Proteobacteria distinguishes rural Himalayan populations from the Americans. **(B)** Heatmap displaying 52 genera with significantly different abundance across the five populations. Bars on the top represent the grouping of individuals in the heatmap columns by their populations or lifestyles. Genera labels in rows are colored by their phylum. Purple: Actinobacteria, dark blue: Bacteroidetes, light red: Elusimicrobia, orange: Firmicutes, light blue: Proteobacteria, magenta: Spirochaetes, light pink: Tenericutes, brown: Verrucomicrobia. Heat map colors reflect relative abundances of each genus.

To characterize the taxonomic differences between populations at a finer level, we repeated the above analysis at the genus level and identified 52 out of 116 genera that showed significant differences in abundance across the five populations (FDR adjusted P-value <0.05, Figure 5B, Supplementary Table 7). Consistent with the differences observed at the phylum level, the rural populations were enriched for several members of Proteobacteria, including *Ruminobacter, Campylobacter, Succinivibrio,* and *Escherichia/Shigella* (Supplementary Figure 7). Among the rural populations, the Chepang foragers were enriched for *Ruminobacter, Campylobacter,* and *Treponema.* Although we did not detect significant differences in abundances of Bacteroidetes across these populations, several members of this phylum distinguished the rural and western populations. The rural Himalayan communities were enriched for *Prevotella, Alloprevotella,* and *Anaerophaga* and significantly depleted in *Bacteroides, Alistipes, Butyricimonas, Odoribacter,* and *Barnesiella*. 29 genera belonging to Firmicutes differed significantly across the five populations and their distribution was complex across these populations (Supplementary Figure 8). Traditional populations were enriched for *Clostridium sensu stricto*, *Catenibacterium, Lactobacillus, Bulleidia*, *Sarcina, Enterococcus, Eubacterium, Oribacterium, Mogibacterium, Mitsuokella, Allisonella, Weissella, Papilbacter* and two unknown genera of Erysipelotrichaceae and Veillonellaceae families. Alternatively, abundances of several *Clostridium* genera, *Oscillibacter, Blautia, Butyriciococcus, Anaerostipes,* and *Flavonifractor* were elevated in the Americans. The Americans also showed highest abundances of *Bifidobacterium* (Actinobacteria) and *Akkermansia* (Verrucomicrobia), both of which were extremely low in the Chepang foragers. Elevated abundances of *Treponema* and *Prevotella* with reduction of *Bacteroides and Bifidobacterium* is a characteristic feature of gut microbiomes of foraging communities [19,24,28,30].

**Figure 8.**
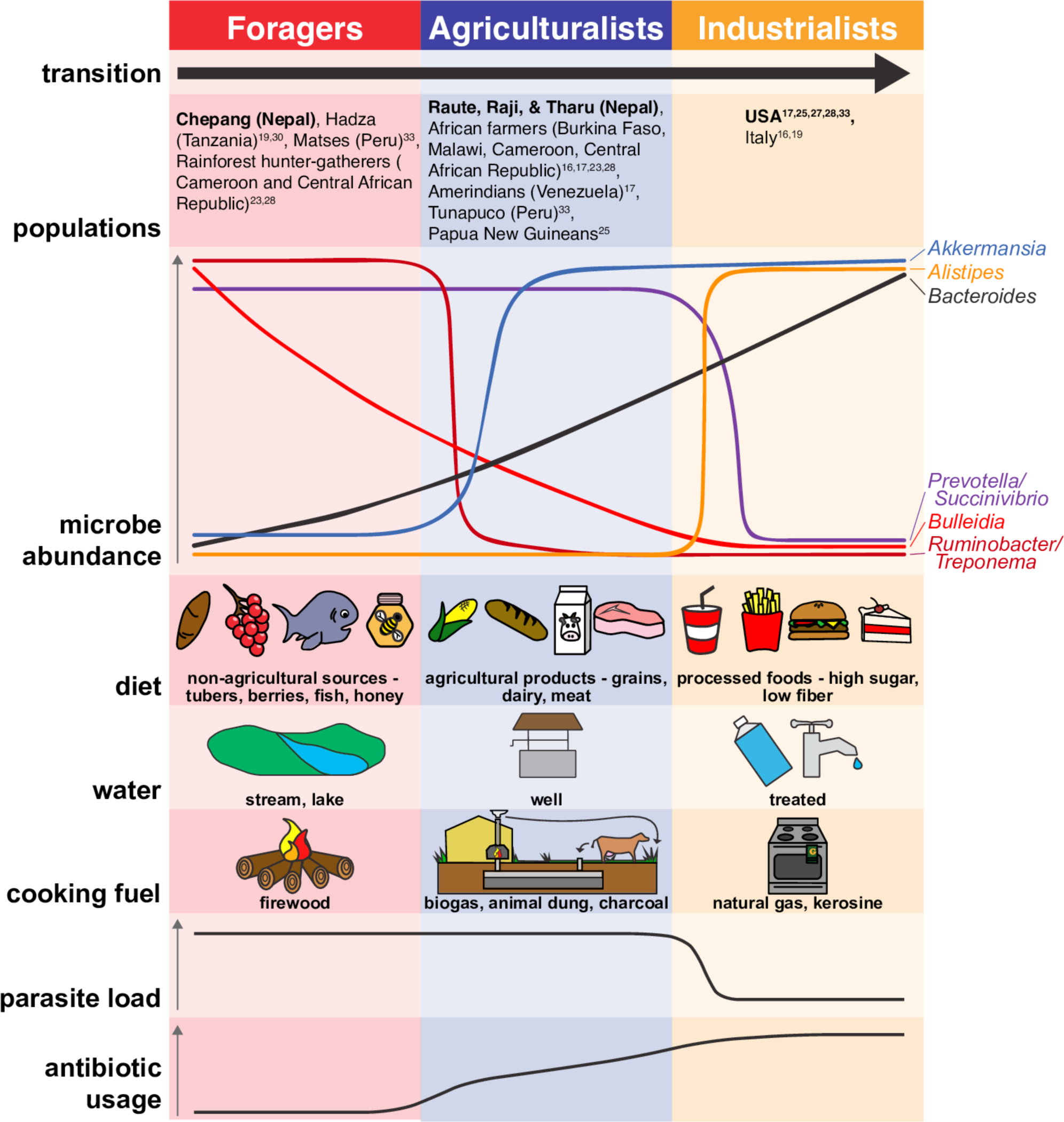
Proposed dynamics of gut microbiome in lifestyle transitions. We propose fluctuations in individual taxa in the gutmicrobiota show complex patterns as humans transition from one lifestyle to another. A few examples of bacterial taxa and their consistent patterns of changes in human populations across the world are shown. Certain genera such as *Treponema* and *Ruminobacter* that are characteristic of hunter-gatherers rapidly decline in agrarians and industrialists. In contrast, taxa such as *Alistipes* and *Akkermansia* rapidly increase in non-foragers. Genera such as *Bacteroides* show a gradual increase from foragers to industrialists and *Bulleidia* show an opposite trend. Higher abundances of taxa such as *Prevotella* and *Succinivibrio* are characteristics of traditionallifestyles and are virtually absent in industrialists. Both dietary and environmental factors are likely to influence the gut microbiome. In this study, source of drinking water was strongly associated with gut microbiome composition. Other environmental factors such as parasite load and antibiotic usage also influence the gut microbiota.

To evaluate whether these taxa reflect lifestyle gradients, we measured the correlations of genus abundances with the coordinates from the PCoA1 axis obtained from the unweighted UniFrac analysis and found strong correlations for 33 of the 52 differentially abundant genera (Spearman’s rho >0.29, q-value <0.05, *correlation test*). *Bacteroides* showed the strongest positive correlation with the PCoA1 (rho=0.78, q-value = 1.9 X 10^-12^, *correlation.test*) while *Ruminobacter, Treponema, Bulleidia,* and *Catenibacterium* showed strong negative correlations (Figure 6), consistent with multiple genera varying with lifestyle differentiation across the five populations.

**Figure 6.**
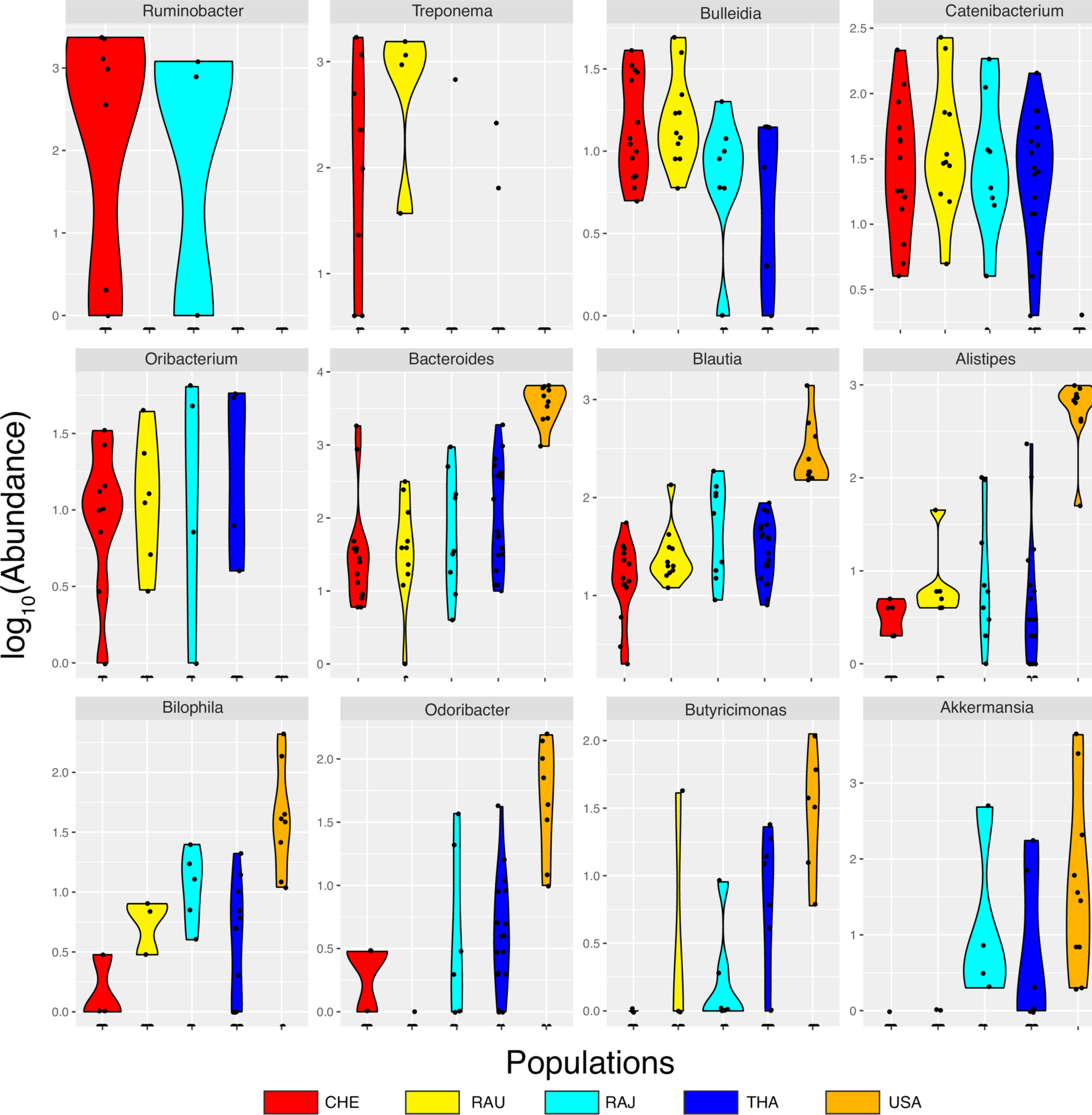
Genera strongly associated with lifestyle gradients. Each of these genera is strongly correlated with the principal axis (PCoA1) of the unweighted UniFrac distances. Violin plots are color coded by populations.

### Factors affecting gut microbiome composition in the Himalaya

We next assessed whether any of the ten dietary and environmental factors that differentiate the Himalayan populations (from Figure 2) correspond to the variation in gut microbiome composition. A canonical correspondence analysis (CCA) revealed that the ten factors collectively explain 38% of the gut microbiome variation within Himalaya while 62% of the variation remained unexplained. Of the ten variables, the source of drinking water and use of solid biomass fuel were significantly associated with the gut microbiome composition in the Himalayan populations (P-value = 0.009 and 0.028 respectively, *ANOVA*), indicating that environmental factors can affect the gut microbiome. Both of these factors contributed most to the first CCA axis (CCA1), which distinguished the Chepang and Raute individuals who drink river water and exclusively burn solid biomass fuel for cooking from the Raji and Tharu who drink underground water and use biogas for cooking (Figure 7). Individuals who drank river water had higher abundances of *Treponema* and those who drank underground water had elevated levels of *Fusobacterium* (q-value<0.05 for both, *Kruskal-Wallis test*). Although cooking fuel was significantly associated at the compositional level, none of the genera reached statistical significance after correcting for multiple testing.

**Figure 7:**
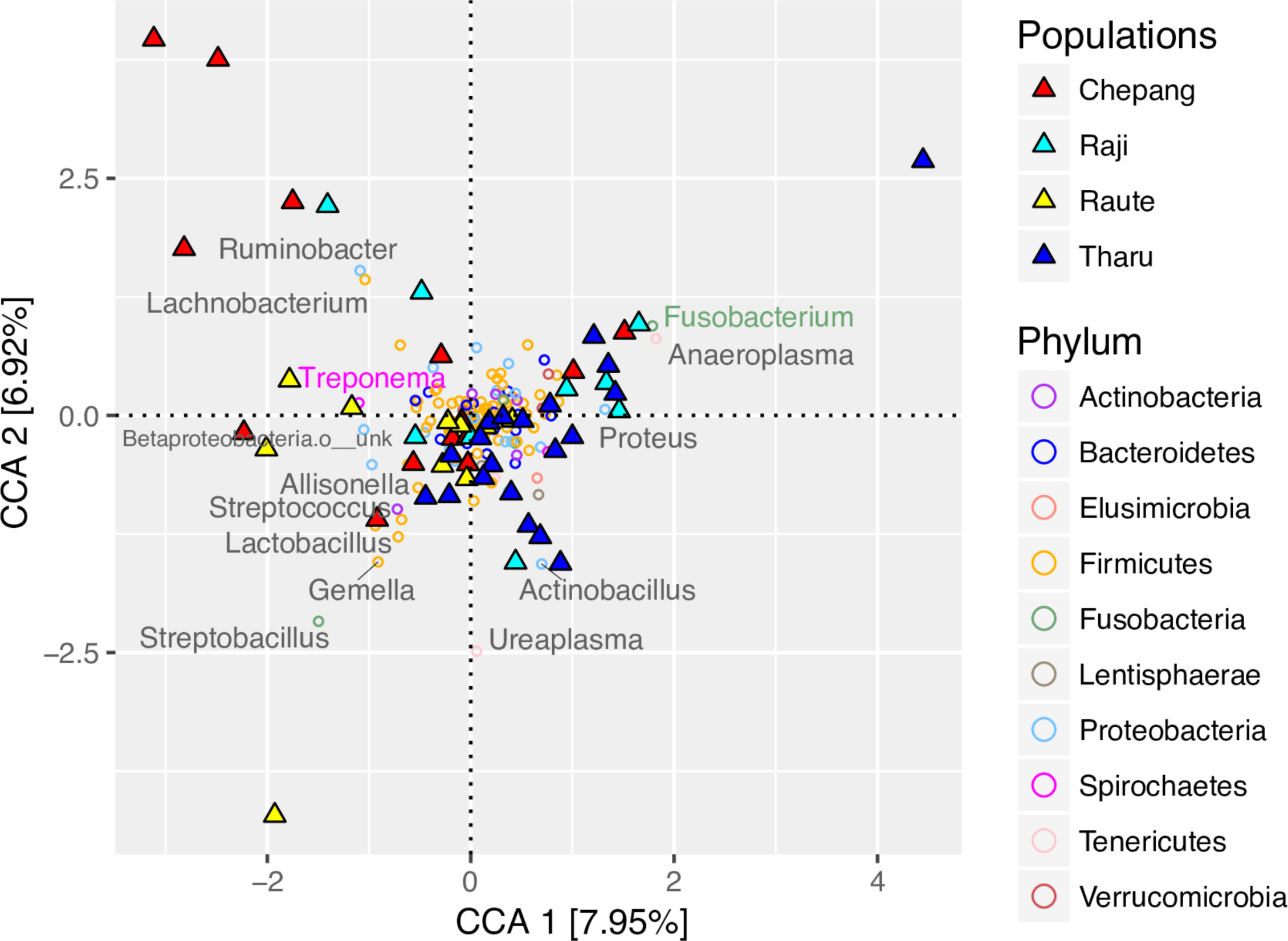
Environmental factors associated with the gut microbiome composition in Himalaya. The two primary CCA axes and the proportion of constrained variance they explain are shown. Triangles represent individuals and circles represent genus. Individuals and genera are color coded by their respective populations and phyla. Drinking water and cooking fuel contributed most to CCA1 and *sisnu* (nettles) contributed most to CCA2. Genera labeled in grey contribute to the top two CCA axes. Among these, *Fusobacterium* and *Treponema* were significantly associated with drinking water.

## DISCUSSION

Several previous reports show that gut microbiomes of traditional populations vary from those of westerners [16,17,19–22,24,25,27–29]. These studies have emphasized that gut bacterial composition differs between traditional and westernized populations, alpha diversity is higher in traditional populations, and diet may be the primary driver of variation in the human gut microbiome. However, because these studies compare human populations that diverged tens of thousands of years ago, it has been difficult to separate the effect of geography and lifestyle on gut microbiome. In this study, we compared the gut microbiome from four rural Himalayan populations with shared ancestries that led nomadic lifestyles until recently and transitioned to farming at various time points in the last three hundred years. Although the individuals in our study have historically cohabited a geographically small region (less than 150K sq. km) in the Himalayan foothills and shared similar diets until recently, their current diets and lifestyles vary. Our results indicate that their gut microbiota strongly mirrors their lifestyles, indicating that the human gut microbiome can undergo pronounced changes within a short time (decades) of departure from foraging (as seen in the Raute and Raji). As dependences on agriculture increases, these changes become more pronounced (as seen in the Tharu). Since these populations have shared ancestries and they cohabit comparable latitudinal regions, such changes in gut microbiota are unlikely to be ascribable to host genetic differences or confounded by geography.

The variations in gut microbiome in the Himalayan populations are consistent with the general patterns observed in many traditional human populations. More importantly, our results suggest certain genera represent conserved gut microbiota markers of human subsistence states (Figure 8). Previous studies of the industrialized microbiota have demonstrated that gut microbiome composition associates with and can be driven by differences in host diet [4–6,8,15,16,22,26,30]. Several genera *Ruminobacter* and *Treponema* that are associated with metabolizing uncultivated plant products and are enriched in the Chepang foragers in this study are also elevated in hunter-gatherers across the world [19,28,30,33]. Moreover, *Prevotella* and *Eubacterium,* which have been previously associated with vegetarian diet in the westerners [5] were enriched in all Himalayan populations relative to Americans. In contrast, taxa associated with animal proteins in diet such as *Bacteroides* and *Blautia* [5,46] were enriched in the Americans relative to Himalayan populations. This is consistent with low animal protein content in diet across Nepal [47].

In addition to diet, environmental factors may also influence the human gut microbiome [7,23,44]. Consistent with these findings, we found that differences in sources of drinking water may exert a detectable effect on the gut microbiota. Differences in mineral and microbial content in drinking water in Nepal has been previously reported [48–51], which may be affecting gut microbiome in our Himalayan participants. Moreover, prolonged exposure to air pollutants have been shown to alter gut microbiome in mice [44]. Whether direct or indirect, breathing polluted air containing higher levels of particulate matters due to solid biomass cooking fuel is linked to gut microbiome composition in our study. In addition, intestinal parasite load has been shown to alter gut microbiota [23]. The association between gut microbiome and parasite load approached significance in our participants as well (P=0.075, *ANOVA*), although it did not reach significance likely due to lower parasite abundance in our participants.

Despite noticeable differences in the gut microbiome composition, we did not observe significant differences in gut bacterial diversity (alpha diversity) across lifestyles in the study populations. Comparisons of populations that reside in similar geographical areas but practice different subsistence strategies such as the BaAka hunter-gatherers and Bantu farmers [28] as well as Matses hunter-gatherers and Tunapuco farmers [33] also showed little differences in alpha diversity. However, these and other traditional populations such as the Hadza [19] have elevated gut bacterial diversity relative to westerners. We did not observe higher alpha diversity in the traditional Himalayan populations relative to the Americans. One possible explanation for this discrepancy could be that latitude is the primary factor that influences gut bacterial diversity. The traditional populations included in previous studies reside in the tropical climate zones, which have higher biodiversity likely affecting both diet and environmental microbial exposures.

In conclusion, our results emphasize the need to study additional traditional populations to understand how geography, climate, diet, and environment affect the gut microbiome. By comparing human populations that reside in a relatively small geographical area, shared a common diet and lifestyle until recently, and are currently practicing different subsistence strategies, we show that human gut microbiome undergoes marked changes within decades of increasing urbanization. Indeed, the extent to which the numerous factors associated with urbanization contribute to gut microbiome change remain to be determined, although gut microbiome extinction events have been shown in experimental models to result from western diet, antibiotics, and chemical laxatives [5–7]. However, the global trends of bacterial taxa within the gut that undergo depletion or enrichment upon lifestyle transitions are striking. The functional consequence of these changes, both in terms of the intrinsic microbial ecology of the gut and the impact on human biology, are critical questions for the field to address. Future work should incorporate metagenomics to characterize the gut microbial variation at finer scales, metabolomics and strain culturing to assess functional differences, and immune and metabolic profiling of these populations. Pursuit of mechanisms by which the gut microbiome interacts with the ecosystems of these populations may reveal conserved connections between microbial and human biology with large implications for industrialized humans who lack these microbes.

## MATERIALS AND METHODS

### Study sites, participating individuals, and sample collection

Stool samples were collected with informed consent from 56 adult participants (over 18 years old) from four indigenous Himalayan populations from Nepal and 10 adult Americans of European descent. Indigenous populations from Nepal included Chepang (N=14), Raji (N=10), Raute (N=12), and Tharu (N=20) inhabiting in Chitwan, Bardia, Dadeldhura, and Sarlahi districts respectively. The samples were collected in winter of 2016 (March and April) with consent from all participants. This work was approved by Ethical Review Board of the Nepal Health Research Council (NHRC) as well as by the Stanford University Institutional Review Board (IRB).

In addition to collecting the fecal samples, we also obtained ethno-linguistic, demographic, environmental, and dietary data from the participants using a survey questionnaire specifically designed for this study. The survey questionnaire assessed participant’s age, gender, diet, health status, use of medication, and behavioral practices such as tobacco and alcohol consumption along with several environmental variables (Supplementary Table 2). In addition, we also visually inspected the stool samples of each individual under the microscope for the presence of intestinal parasites (triplicate slides per individual). Participants’ responses to survey data questionnaires are included in Supplementary Table 4.

### DNA extractions

Freshly produced stool samples from the Himalayan participants were collected on a clean OMNIgene gut accessory collection paper (OM-AC1). About 500mg of the stool samples was transferred to the OMNIgene gut kit collection tube containing the stabilizing buffer using the clean spatula provided with the kit. The tubes were shaken hard in a back and forth motion until the fecal samples were completely homogenized. Tubes were transported at room temperature within 48-72 hours of collection to Tribhuvan University Institute of Medicine, Kathmandu, Nepal where they were transferred to -80°C until DNA extraction. DNA was extracted using MolBio Power Soil Kit according to the manufacturer’s protocol. Extracted DNA was shipped to Stanford University on dry ice and stored at -20°C until sequencing. Samples from Americans were collected from volunteers at Stanford University in a 15ml centrifuge tubes and transported to the laboratory on ice. Half of each sample was immediately frozen at -80°C. From the other half, 500mg stool was transferred to OMNIgene collection tubes and kept at room temperature for 48-72 hours after which they were stored at -80°C. DNA was extracted from both sets of samples simultaneously using MolBio Power Soil Kit according to the manufacturer’s protocol and stored at -20°C until sequencing.

### 16S sequencing and analyses

The V4 region of the 16S rRNA gene was PCR amplified using the primers and protocols described previously [52]. The amplified DNA fragments were multiplexed and subjected to paired-end sequencing using Illumina MiSeq. Of the 66 samples, one yielded very low levels of DNA and another failed the paired end sequencing. After discarding these two samples, the final dataset included 64 individuals (14 Chepang, 9 Raji, 11 Raute, 20 Tharu, and 10 Americans). The amplification primers and barcodes used for multiplexing are described in Supplementary Table 4.

Paired-end reads were processed using DADA2 [53] and subsequently analyzed in R using *phyloseq* [54]. In order to identify high quality sequences, reads were trimmed to 150 bp. Sequences with N nucleotides and/or >2 expected errors were discarded (maxN=0, maxEE=2, truncQ=2) and sequence variants were inferred by pooling reads from all samples (pool=TRUE). Sequence tables were then created by merging paired-end reads. A naïve Bayesian classifier method [55] implemented in DADA2 algorithm was used to assign taxonomy using the RDP v14 training set [56]. Multiple alignment was conducted using DECIPHER [57] package in R and a maximum likelihood phylogenetic tree was constructed using *phangorn* [58] with a neighbor-joining tree as the starting point.

A total of 1,183,760 merged reads passed quality control and 1630 taxa were initially identified. After removing chimeric sequences, which constituted 22% of the reads, 921,345 merged reads remained. Further elimination of low abundance phyla – Synergistetes and Deferribacteres – that were observed only once across all samples resulted in 883 taxa in the dataset. After quality control, mean (±SD) sequencing depth per sample was 11570 (±4653). We performed three technical replicates of the frozen sample for one individual and a total of five replicates for two additional individuals for the OMNI samples. Since we did not observe marked differences in the technical replicates (Supplementary Figure 3), we retained the sample with highest coverage for these individuals. After removing the replicate samples, 64 individuals and 875 taxa remained in the final dataset.

### Random forest classifier model

One hundred random forest classifiers (RFC) with 50 to 5000 trees were constructed using all 35 variables (Supplementary Table 3) from the survey data using ‘randomForest’ R package [59]. We also repeated this analysis on the 16S data and reported the RFC with smallest out of bag error rate for both analyses.

### Statistical analyses

Correspondence analysis of the survey data was performed using FactoMineR package in R [60]. Canonical correspondence analysis was performed at the genus level (taxa collapsed based on genus names) by calling functions from *vegan* package via *phyloseq*. Phylogenetic diversity was computed by rarefying the samples to various depths starting from 500-3000 sequences per sample. Alpha diversity was measured using species richness, Shannon’s H, Simpson’s D, and Fisher’s alpha, calculated as the mean values from 100 iterations at each depth. Kruskal-Wallis tests were used to assess the significance of differences in each of the alpha diversity metrics between populations at each rarefaction depth. Differences in rarefaction depth did not alter significance of the observed differences. Hence, we chose to report results from rarefaction depth of 3000, which was the maximum depth that allowed inclusion of all of the samples. Beta diversity was assessed using Bray-Curtis as well as unweighted and weighted UniFrac distances calculated by log transformation of the non-rarefied 16S count data. Permutational multivariate analysis of variance (PERMANOVA) was performed using the *vegan* package in R [61]. For all PERMANOVA analyses, 10000 randomizations were performed to assess the statistical significance. In order to identify differentially abundant taxa at the phylum and genus levels, we first agglomerated the taxa abundance (counts) at each taxonomic level respectively. The differences in taxa abundance (counts) were then assessed using the DESeq2 package [45]. Multiple testing corrections were performed by computing false discovery rates (FDR) using Benjamini-Hochenberg method and adjusted p-values < 0.05 were considered statistically significant.

## AUTHOR CONTRIBUTIONS

Conceived by A.R.J, E.R.D, J.S, and C.D.B; sample collection, DNA extraction, and sequencing were performed by A.R.J, Y.G, G.P.G, D.B, S.T, and K.N; parasite characterization by D.B and S.T under the supervision of J.B.S; data analysis was conducted by A.R.J and E.R.D, and under the supervision of S.H, J.S and C.D.B. Resources were provided by Y.G, G.P.G, J.B.S, J.S, and C.D.B. Manuscript prepared by A.R.J with input from all authors.

## ACKNOWLEDGEMENTS

We would like to thank the Nepal Health Research Council (NHRC) under the Government of Nepal Ministry of Health for providing research permits to conduct our work in Nepal. We are grateful to Mr. Biswash Chepang for community outreach and participant recruitment in the Chepang community. We express our gratitude towards all the participants in this study. This work was partly supported by Center for Human and Evolutionary Genomics (CEHG) Seed Award to A.R.J., and grants from National Institutes of Health R01-DK085025 and DP1-AT00989201 to J.L.S. E.R.D is supported by NIH F32 DK109595. C.D.B and J.L.S are Chan Zuckerberg Biohub and Chan Zuckerberg Biohub Microbiome investigators.

## SUPPORTING INFORMATION

**Supplementary Figure 1: Dietary and environmental factors associated with lifestyle gradients in the Himalaya.** Several dietary factors distinguished the four Himalayan populations included in this study. Foraged plants such as *sisnu* (nettles) and *jaand*, a slushy alcoholic beverage made from fermenting millet or corn, are staples of the Chepang diet. Although not recorded in the survey data, our Chepang participants reported that due to lack of irrigation, they are unable to grow rice and are limited to growing crops that require less water such as buckwheat, millet, and corn and forage for tubers (g*ittha vyakur*) in the forest. In contrast, alcohol use was minimal among the Raute, Raji, and Tharu. Moreover, perceived food scarcity was higher in the Chepang and Raute, both of which reside in remote villages relative to the Raji and Tharu. Although meat consumption was low across all four populations, the Tharu consumed animal products such as yogurt most frequently. According to our Tharu participants, *ghonghi* (snails) are staples in their diet, although this dietary parameter was not included in our survey. In addition to diet, several environmental factors also differed across the Himalayan populations. The Chepang and Raute who reside in remote villages still fetch their drinking water from rivers and streams. Conversely, Raji and Tharu who reside in more urbanized areas have installed tube wells in their homes enabling access to underground water for drinking. The use of solid biomass fuel (SBM) was lower in the Tharu and Raji as they frequently used non-solid biomass fuel (NSBM) such as biogas. Conversely, the Chepang and Raute are still completely dependent on burning firewood for cooking. Although, we detect low overall levels of intestinal parasites in our participants Ascaris, Entamoeba, Trichuris, Hymenolepis, and Coccidia appear in some individuals. Parasite infection was highest in the Chepang, intermediate in the Raute and Raji, and lowest in the Tharu. Smoking and tobacco consumption was higher in the Tharu and Chepang relative to Raji and Raute.

**Supplementary Figure 2: 16S sequencing and quality filtering. (A)** Sequencing depth for each taxa and each sample before filtering. Over 1600 read sequence variants (RSVs) were initially identified but many were chimeric and detected by a single read. **(B)** Removal of chimera did not reduce sequencing depth for the taxa or for the samples. **(C)** Abundance of the phyla in the dataset. Deferribacteres and Synergistes were detected in only a few individuals and were lowly abundant (read count < 4) and were removed. **(D)** After quality filtering of chimera and low abundance taxa, twelve phyla and a total of 883 taxa remained in the dataset.

**Supplementary Figure 3: Comparison of frozen and OMNIgene samples.** Since flash freezing of the samples was not possible in the remote sampling areas, we used commercially available DNAgenotek OMNIgene kits to collect stool samples from the four Himalayan populations. We also collected stool samples from 10 Americans of European descent from Palo Alto. We divided these samples into two sets, the first set was trasferred into OMNIgene kits and the second set was frozen at -80C. The OMNIgene kits containing the stool samples were kept at room temperatures for 24-72 hours then they were frozen at -80C. DNA extraction, 16S amplification (V4), and sequencing was performed simultaneously for both sets of samples. This allowed us to determine whether the kit collections in the field could faithfully reproduce expected microbiome profiles as well as freshly frozen stool. **(A)** Analysis of gut bacterial community using Principal Coordinate Analysis (PCoA) of unweighted and weighted UniFrac distances showed no significant differences between the sampling methods (P>0.05 for both distances, *PERMANOVA*). Replicate samples from the same individual also tended to be in close proximity of one another in both analyses. **(B)** Alpha diversity assessed using species richness, Fisher index, and Shannon index was not significantly different between the two methods (P>0.05, *Kruskal-Wallis test*). **(C)** Although comparison of frozen and OMNI samples showed little differences, abundance of Euryarcheota and Cyanobacteria/chroloplast were lower and higher in OMNI samples, respectively. Both constituted negligible fractions of gut bacteria and were removed from further analyses. **(D)** Comparison of differences in taxa abundances at the genus level using a negative binomial generalized linear model for differential abundance analysis as implemented in DESeq2 demonstrated that none of the genera differed significantly between the sampling methods (FDR adjusted P-values >0.05). Hence, these results collectively demonstrate that sampling using OMNIgene kits did not introduce major biases in our data.

**Supplementary Figure 4: Differences in gut microbiome compositions across lifestyles. (A)** Visualizationusing a PCoA of the Bray-Curtis (left) and weighted UniFrac distances (right). Each dot represents an individual and colors indicate the populations. In both analyses, PCoA1 explains most variation in the dataset (21.5% and 34.5% of the Bray-Curtis and weighted UniFrac distances respectively). **(B)** Distribution of populations along the PCoA1 axis show patterns of separation by lifestyles. Chepang foragers (red), Raute (yellow) and Raji (cyan), Tharu farmers (blue), and Americans (orange). **(C)** In both cases, PCoA1 was strongly correlated with CA1 obtained from the analysis of the survey data. Spearman’s rho for Bray-Curtis and weighted UniFrac were 0.47 and 0.28 respectively (P<0.05 for both, *correlation test*). Correlations between CA2 and PCoA1 were insignificant (P>0.05 for both distances, *correlation test*).

**Supplementary Figure 5: Visualization of distinctions in gut microbial communities across population using PCoA.** PCoA of the unweighted and weighted UniFrac distances (top and middle respectively) and Bray-Curtis distance (bottom). All four plots on each row differ only in coloring of the dots to help visualize the distribution of individuals in each population.

**Supplementary Figure 6: Variation in gut microbiota within Himalaya.** Columns show PCoA of the three distance matrices of the four Himalayan populations after removing Americans from the analysis. Top row shows the top two PCoA axes and variance explained. Significant differences in gut microbiome composition within Himalaya was observed for all three distances (P<0.05, *PERMANOVA*). Bottom row shows the distribution of the Himalayan populations along the PCoA 1 axis. The separation between Chepang foragers (red) and Tharu farmers (blue) is the strongest within Himalaya with the two transitioning Raute and Raji populations as intermediates.

**Supplementary Figure 7: Abundances of differentially abundant genera across populations.** Each subplot shows abundance of an individual taxa in the five populations. Differentially abundant genera from Bacteroidetes (A), Proteobacteria (B), Verrucomicrobia (C), Spirocheates (D), Actinobacteria (E), Elusimicrobia (F), and Tenericutes (G). Labels with “c__unk” and “f__unk” indicate taxa with unknown Class and Family respectively.

**Supplementary Figure 8: Complex patterns of differential abundances of Firmicutes across populations.** Several genera are significantly enriched in the rural Himalayan populations and others are depleted. Labels with “g__unk” and “o__unk” indicate taxa with unknown genus and order respectively.

**Supplementary Table 1: Populations, their subsistence strategies, and sample sizes:** For the 10 Americans we compared frozen samples to those collected using OMNIgene collection kits. We also performed 3 technical replicate sequencing for 2 Americans. Although we attempted to balance the numbers of males and females from each population there were slightly higher representation of females than males in this study.

**Supplementary Table 2: Survey questionnaire:** Survey data were collected for the 53 of the 54 individuals from the four Himalayan populations. One individual consented to donating samples but was not interested in participating in the survey. We included the sample and removed this individual from the survey data analyses. Prolonged exposure to pollutants generated during combustion of solid biomass fuel such as firewood or animal dungs due to indoor cooking has the potential to alter gut microbiome. Hence, we assessed the fuel types used for cooking and location of kitchen in our Himalayan participants. We also inquired about the sources of drinking water among our participants. None of the participants filtered or purified water before drinking. Thus this variable was excluded from analysis. We surveyed three replicates from each the stool samples under a microscope to identify parasites Ascaris, Entamoeba, Trichuris, Hymenolepis, and Coccidia. If any of these parasites were present, the individuals were labeled positive. Frequency of plant and animal products in diet were also recorded. Binary responses were coded as 0 and 3, frequency variables were coded as 0,1,2,3 for least frequent to most frequent.

**Supplementary Table 3: Random forest classifications:** Summary of random forest classification using survey data (A) and 16S OTU table (B). Lowest out of bag error (3%) for the survey data was obtained with 2750 trees and lowest out of bag error (32%) for the 16S data was obtained with 1950 trees.

**Supplementary Table 4: Primers, sequencing depth, and survey data:** Amplification primers, barcodes used for multiplexing and sequencing depth for samples in this study along with survey data collected from participants. “NA” indicates missing data.

**Supplementary Table 5: Mean distances within and between populations:** Average pairwise distances between individuals within and between populations computed using Bray-Curtis, unweighted UniFrac, and weighted UniFrac matrices.

**Supplementary Table 6: Significantly different phyla across populations:** Summary table of differential abundance of phyla (taxa collapsed based on phylum names) across the five populations. Statistical significance was assessed using a negative binomial generalized linear model as implemented in DESeq2. Multiple testing corrections were performed by computing false discovery rates (FDR) using Benjamini-Hochenberg method and multiple testing adjusted P-values < 0.05 were considered statistically significant.

**Supplementary Table 7: Significantly different genera across populations:** Summary table of differential abundance of genera (taxa collapsed based on genus names) across the five populations. Statistical significance was assessed using a negative binomial generalized linear model as implemented in DESeq2. Multiple testing corrections were performed by computing false discovery rates (FDR) using Benjamini-Hochenberg method and multiple testing adjusted P-values < 0.05 were considered statistically significant.

## REFERENCES

[1] Nicholson JK, Holmes E, Kinross J, Burcelin R, Gibson G, Jia W, et al. Host-gut microbiota metabolic interactions. Science (80-). 2012; doi:10.1126/science.1223813

[2] Belkaid Y, Hand TW. Role of the microbiota in immunity and inflammation. Cell. 2014. doi:10.1016/j.cell.2014.03.011

[3] van Nood E, Vrieze A, Nieuwdorp M, Fuentes S, Zoetendal EG, de Vos WM, et al. Duodenal Infusion of Donor Feces for Recurrent Clostridium difficile. N Engl J Med. 2013; 368: 407–415. doi:10.1056/NEJMoa1205037

[4] Wu GD, Chen J, Hoffmann C, Bittinger K, Chen Y, Keilbaugh SA, et al. Linking Long-Tem Dietary Patterns with Gut Microbial Enterotypes. Science (80-). 2011;334: 105–109. doi:10.1126/science.1208344

[5] David LA, Maurice CF, Carmody RN, Gootenberg DB, Button JE, Wolfe BE, et al. Diet rapidly and reproducibly alters the human gut microbiome. Nature. 2013; doi:10.1038/nature12820

[6] Sonnenburg ED, Smits SA, Tikhonov M, Higginbottom SK, Wingreen NS, Sonnenburg JL. Diet-induced extinctions in the gut microbiota compound over generations. Nature. 2016;529: 212–215. doi:10.1038/nature16504

[7] Dethlefsen L, Huse S, Sogin ML, Relman DA. The pervasive effects of an antibiotic on the human gut microbiota, as revealed by deep 16s rRNA sequencing. PLoS Biol. 2008;6: 2383–2400. doi:10.1371/journal.pbio.0060280

[8] Ji Y, Sun S, Goodrich JK, Kim H, Poole AC, Duhamel GE, et al. Diet-induced alterations in gut microflora contribute to lethal pulmonary damage in TLR2/TLR4-deficient mice. Cell Rep. 2014;8: 137–149. doi:10.1016/j.celrep.2014.05.040

[9] Turnbaugh PJ, Hamady M, Yatsunenko T, Cantarel BL, Duncan A, Ley RE, et al. A core gut microbiome in obese and lean twins. Nature. 2009;457: 480–484. doi:10.1038/nature07540

[10] Ley RE, Backhed F, Turnbaugh P, Lozupone CA, Knight RD, Gordon JI. Obesity alters gut microbial ecology. Proc Natl Acad Sci. 2005;102: 11070–11075. doi:10.1073/pnas.0504978102

[11] Ley RE, Turnbaugh PJ, Klein S, Gordon JI. Microbial ecology: human gut microbes associated with obesity. [Internet]. Nature. 2006. doi:10.1038/4441022a

[12] Shreiner AB, Kao JY, Young VB. The gut microbiome in health and in disease. Curr Opin Gastroenterol. 2015;31: 69–75. doi:10.1097/MOG.0000000000000139

[13] Goodrich JK, Waters JL, Poole AC, Sutter JL, Koren O, Blekhman R, et al. Human genetics shape the gut microbiome. Cell. 2014; doi:10.1016/j.cell.2014.09.053

[14] Shreiner AB, Kao JY, Young VB. The gut microbiome in health and in disease. Curr Opin Gastroenterol. 2016;31: 69–75. doi:10.1097/MOG.0000000000000139.The

[15] Sonnenburg ED, Sonnenburg JL. Starving our microbial self: The deleterious consequences of a diet deficient in microbiota-accessible carbohydrates. Cell Metabolism. 2014. pp. 779–786. doi:10.1016/j.cmet.2014.07.003

[16] De Filippo C, Cavalieri D, Di Paola M, Ramazzotti M, Poullet JB, Massart S, et al. Impact of diet in shaping gut microbiota revealed by a comparative study in children from Europe and rural Africa. Proc Natl Acad Sci. 2010; doi:10.1073/pnas.1005963107

[17] Yatsunenko T, Rey FE, Manary MJ, Trehan I, Dominguez-Bello MG, Contreras M, et al. Human gut microbiome viewed across age and geography. Nature. 2012;486: 222–227. doi:10.1038/nature11053

[18] Tyakht A V., Kostryukova ES, Popenko AS, Belenikin MS, Pavlenko A V., Larin AK, et al. Human gut microbiota community structures in urban and rural populations in Russia. Nat Commun. 2013; doi:10.1038/ncomms3469

[19] Schnorr SL, Candela M, Rampelli S, Centanni M, Consolandi C, Basaglia G, et al. Gut microbiome of the Hadza hunter-gatherers. Nat Commun. Nature Publishing Group; 2014;5: 3654. doi:10.1038/ncomms4654

[20] Moeller AH, Li Y, Mpoudi Ngole E, Ahuka-Mundeke S, Lonsdorf E V., Pusey AE, et al. Rapid changes in the gut microbiome during human evolution. Proc Natl Acad Sci. 2014;111: 16431–16435. doi:10.1073/pnas.1419136111

[21] Escobar JS, Klotz B, Valdes BE, Agudelo GM. The gut microbiota of Colombians differs from that of Americans, Europeans and Asians. BMC Microbiol. 2014; doi:10.1186/s12866-014-0311-6

[22] O’Keefe SJD, Li J V., Lahti L, Ou J, Carbonero F, Mohammed K, et al. Fat, fibre and cancer risk in African Americans and rural Africans. Nat Commun. 2015;6: 6342. doi:10.1038/ncomms7342

[23] Morton ER, Lynch J, Froment A, Lafosse S, Heyer E, Przeworski M, et al. Variation in Rural African Gut Microbiota Is Strongly Correlated with Colonization by Entamoeba and Subsistence. PLoS Genet. 2015;11: 1–28. doi:10.1371/journal.pgen.1005658

[24] Obregon-Tito AJ, Tito RY, Metcalf J, Sankaranarayanan K, Clemente JC, Ursell LK, et al. Subsistence strategies in traditional societies distinguish gut microbiomes. Nat Commun. 2015;6: 6505. doi:10.1038/ncomms7505

[25] Iné Martínez A, Stegen JC, Greenhill AR, Walter J, Martínez I, Maldonado-Gó mez MX, et al. The Gut Microbiota of Rural Papua New Guineans: Composition, Diversity Patterns, and Ecological Processes. Cell Rep. 2015; doi:10.1016/j.celrep.2015.03.049

[26] Zhang J, Guo Z, Lim AAQ, Zheng Y, Koh EY, Ho D, et al. Mongolians core gut microbiota and its correlation with seasonal dietary changes. Sci Rep. 2015; doi:10.1038/srep05001

[27] Clemente JC, Pehrsson EC, Blaser MJ, Sandhu K, Gao Z, Wang B, et al. The microbiome of uncontacted Amerindians. Sci Adv. 2015;1: e1500183–e1500183. doi:10.1126/sciadv.1500183

[28] Gomez A, Petrzelkova KJ, Burns MB, Yeoman CJ, Amato KR, Vlckova K, et al. Gut Microbiome of Coexisting BaAka Pygmies and Bantu Reflects Gradients of Traditional Subsistence Patterns. Cell Rep. 2016;14: 2142–2153. doi:10.1016/j.celrep.2016.02.013

[29] Dehingia M, Thangjam devi K, Talukdar NC, Talukdar R, Reddy N, Mande SS, et al. Gut bacterial diversity of the tribes of India and comparison with the worldwide data. Sci Rep. 2016; doi:10.1038/srep18563

[30] Smits SA, Leach J, Sonnenburg ED, Gonzalez CG, Lichtman JS, Reid G, et al. Seasonal cycling in the gut microbiome of the Hadza hunter-gatherers of Tanzania. Science (80-). 2017;357: 802–806. doi:10.1126/science.aan4834

[31] Dikongué E, Ségurel L. Latitude as a co-driver of human gut microbial diversity? BioEssays. 2017;39. doi:10.1002/bies.201600145

[32] Creanza N, Kolodny O, Feldman MW. Cultural evolutionary theory: How culture evolves and why it matters. Proc Natl Acad Sci. 2017;114: 7782–7789. doi:10.1073/pnas.1620732114

[33] Obregon-Tito AJ, Tito RY, Metcalf J, Sankaranarayanan K, Clemente JC, Ursell LK, et al. Subsistence strategies in traditional societies distinguish gut microbiomes. Nat Commun. Nature Publishing Group; 2015;6: 6505. doi:10.1038/ncomms7505

[34] Fortier J, Rastogi K. Sister Languages? Comparative Phonology of Two Himalayan Languages. Nepal Linguist. 2004;21: 42–52.

[35] van Driem G. Languages of the Himalayas I. Brill; 2002.

[36] Chaubey G, Singh M, Crivellaro F, Tamang R, Nandan A, Singh K, et al. Unravelling the distinct strains of Tharu ancestry. Eur J Hum Genet. 2014; 1–9. doi:10.1038/ejhg.2014.36

[37] Reinhard J. The Raute: Notes on a Nomadic Hunting and Gathering Tribe of Nepal. Kailash, A J Himal Stud. 1974;2: 233–271.

[38] Valli E. Golden Harvest of the Raji. National Geographic. 1998: 85–105.

[39] Gurung GM. Economic Modernization in a Chepang Village in Nepal. Occas Pap Sociol Anthropol. 1990;2: 32–39.

[40] Dahal DR. Economic Development through Indigenous Means: A Case of Indian Migration in the Nepal Terai. Contrib Nepalese Stud. 1983;11.

[41] Fortier J. The Ethnography of South Asian Foragers. Annu Rev Anthropol. 2009;38: 99–114. doi:10.1146/annurev-anthro-091908-164345

[42] Hodgson B. On the Chepang and Kusunda Tribes of Nepál. Essays on the Languages, Literature, and Religion of Nepál and Tibet. First. Cambridge: Cambridge University Press, Cambridge Library Collection. Web. 15 October 2015.; 1848. pp. 191–200. doi:http://dx.doi.org/10.1017/CBO9781139507387.016

[43] Fullerton DG, Bruce N, Gordon SB. Indoor air pollution from biomass fuel smoke is a major health concern in the developing world. Transactions of the Royal Society of Tropical Medicine and Hygiene. 2008. pp. 843–851. doi:10.1016/j.trstmh.2008.05.028

[44] Kish L, Hotte N, Kaplan GG, Vincent R, Tso R, Gänzle M, et al. Environmental Particulate Matter Induces Murine Intestinal Inflammatory Responses and Alters the Gut Microbiome. PLoS One. 2013;8. doi:10.1371/journal.pone.0062220

[45] Anders S, Huber W. Differential expression analysis for sequence count data. EMBL, Heidelberg, Ger. 2010; doi:10.1186/gb-2010-11-10-r106

[46] Claesson MJ, Jeffery IB, Conde S, Power SE, O’connor EM, Cusack S, et al. Gut microbiota composition correlates with diet and health in the elderly. Nature. 2012;488: 178–184. doi:10.1038/nature11319

[47] Aryal KK, Mehata S, Neupane S, Vaidya A, Dhimal M, Dhakal P, et al. The burden and determinants of non communicable diseases risk factors in Nepal: Findings from a nationwide STEPS survey. PLoS One. Public Library of Science; 2015;10.

[48] Singh A, Smith LS, Shrestha S, Maden N. Efficacy of arsenic filtration by Kanchan Arsenic Filter in Nepal. J Water Health. 2014; doi:10.2166/wh.2014.148

[49] Ghaju Shrestha R, Tanaka Y, Malla B, Bhandari D, Tandukar S, Inoue D, et al. Next-generation sequencing identification of pathogenic bacterial genes and their relationship with fecal indicator bacteria in different water sources in the Kathmandu Valley, Nepal. Sci Total Environ. 2017; doi:10.1016/j.scitotenv.2017.05.105

[50] WaterAid. Groundwater Quality : Nepal. British Geological Survey. 2001.

[51] Inoue D, Hinoura T, Suzuki N, Pang J, Malla R, Shrestha S, et al. High-Throughput DNA Microarray Detection of Pathogenic Bacteria in Shallow Well Groundwater in the Kathmandu Valley, Nepal. Curr Microbiol. 2014; doi:10.1007/s00284-014-0681-x

[52] Caporaso JG, Lauber CL, Walters W a, Berg-Lyons D, Huntley J, Fierer N, et al. Ultra-high-throughput microbial community analysis on the Illumina HiSeq and MiSeq platforms. ISME J. 2012;6: 1621–1624. doi:10.1038/ismej.2012.8

[53] Callahan BJ, McMurdie PJ, Rosen MJ, Han AW, Johnson AJA, Holmes SP. DADA2: High-resolution sample inference from Illumina amplicon data. Nat Methods. 2016;13: 581–583. doi:10.1038/nmeth.3869

[54] McMurdie PJ, Holmes S. Phyloseq: An R Package for Reproducible Interactive Analysis and Graphics of Microbiome Census Data. PLoS One. 2013; doi:10.1371/journal.pone.0061217

[55] Wang Q, Garrity GM, Tiedje JM, Cole JR. Naiive Bayesian classifier for rapid assignment of rRNA sequences into the new bacterial taxonomy. Appl Environ Microbiol. 2007; doi:10.1128/AEM.00062-07

[56] Cole JR, Wang Q, Cardenas E, Fish J, Chai B, Farris RJ, et al. The Ribosomal Database Project: Improved alignments and new tools for rRNA analysis. Nucleic Acids Res. 2009; doi:10.1093/nar/gkn879

[57] Wright ES. DECIPHER: harnessing local sequence context to improve protein multiple sequence alignment. BMC Bioinformatics. 2015; doi:10.1186/s12859-015-0749-z

[58] Schliep KP. phangorn: Phylogenetic analysis in R. Bioinformatics. 2011; doi:10.1093/bioinformatics/btq706

[59] Liaw A, Wiener M. Breiman and Cutler’s Random Forests for Classification and Regression [Internet]. 2015.

[60] Lê S, Josse J, Husson F. FactoMineR : An R Package for Multivariate Analysis. J Stat Softw. 2008;25: 253–258. doi:10.18637/jss.v025.i01

[61] Oksanen J, Blanchet FG, Kindt R, Legendre P, Minchin PR, O’Hara RB, et al. Package “vegan.” R Packag ver 20–8. 2013; 254. doi:10.4135/9781412971874.n145

